# RNAseq in the mosquito maxillary palp: a little antennal RNA goes a long way

**DOI:** 10.1101/016998

**Authors:** David C. Rinker, Xiaofan Zhou, Ronald Jason Pitts, Patrick L. Jones, Antonis Rokas, Laurence J Zwiebel

## Abstract

A comparative transcriptomic study of mosquito olfactory tissues recently published in *BMC Genomics* (Hodges et al, 2014) reported several novel findings that have broad implications for the field of insect olfaction. In this brief commentary, we outline why the conclusions of Hodges et al. are problematic under the current models of insect olfaction and then contrast their findings with those of other RNAseq based studies of mosquito olfactory tissues. We also generated a new RNAseq data set from the maxillary palp of *Anopheles gambiae* in an effort to replicate the novel results of Hodges et al. but were unable to reproduce their results. Instead, our new RNAseq data support the more straightforward explanation that the novel findings of Hodges et al. were a consequence of contamination by antennal RNA. In summary, we find strong evidence to suggest that the conclusions of Hodges et al were spurious, and that at least some of their RNAseq data sets were irrevocably compromised by cross-contamination between samples.

## INTRODUCTION

A recent RNAseq study in *BMC Genomics* [1], reported the presence of an unexpectedly large variety (~50) of odorant receptors (ORs) in the maxillary palps of both *Anopheles coluzzii* (formerly known as *An. gambiae,* M form) and *An. quadriannulatus*. As a result of these data, Hodges et al. concluded that the functional role of the mosquito maxillary palp should be recast from one of relatively conserved, olfactory simplicity into the “more expansive role” of a much more complex chemosensory appendage. If correct, Hodges et al.’s conclusions would overturn a large body of literature on the electrophysiology and molecular composition of the antennae and maxillary palps of insects, particularly as they pertain to *Drosophila* and mosquitoes. In that light, we felt that a careful scrutiny of the data presented in Hodges et al. was warranted.

Our current understanding of the olfactory role of the mosquito maxillary palp is based in large part upon its morphological, electrophysiological and molecular characteristics, many of which it shares with other files such as *Drosophila*. In contrast to the antenna, the maxillary palps of Diptera appear to be chemosensory appendages of limited odor coding complexity, evidenced by a small variety of palp-specific chemoreceptors that are housed in a relatively small number of (~60) morphologically identical chemosensory sensilla [2-4]. Specifically, in the malaria vector mosquito *An. coluzzii*, Lu et al. comprehensively characterized the spatial organization, electrophysiology and OR-composition of the sensilla of the maxillary palp [2]. We observed a uniform class of capitate peg sensilla that was innervated by three stereotypic olfactory receptor neurons (ORNs) and that displayed identical electrophysiological response profiles to odorant stimuli. Each of the two smaller ORNs showed evidence for only two tuning (i.e., odorant specifying) ORs (AgOr8 and AgOr28) while the third, larger neuron showed no evidence for the presence of any ORs (confirmed by the lack of the ubiquitous OR coreceptor, Orco) and expressed instead the three heteromultimeric gustatory receptors (AgGR22,23,24) that are responsible for the detection of CO_2_, an olfactory sensitivity long considered to be a physiological hallmark of the mosquito maxillary palp [5]. These patterns of morphologic simplicity and physiologic uniformity appear conserved across a wide range of mosquito taxa [2, 6-11] and are highly analogous to the simplicity and uniformity present in the maxillary palp of *Drosophila.*

The findings of Hodges et al. challenge these paradigms in three major ways. First, most of the newly elucidated palpal tuning ORs reported by Hodges et al. are also present in the antenna, with several of them being among the most highly expressed ORs in both tissues. This is in direct opposition to prior observations showing that OR expression in Diptera possess tissue specific patterns of zonal expression that segregate in a mutually exclusive fashion between the maxillary palp and antenna [6, 8, 11-14], with several of the key regulatory underpinnings of this segregation having already been identified in *D. melanogaster* [13]. Second, the ~50 novel palpal ORs reported by Hodges et al. in *An. coluzzii* maxillary palps present a seemingly intractable problem of localization as individual ORNs in flies are considered to generally express only one type of tuning OR [15-17]. Given that the thorough characterization of palpal ORNs by Lu et al. consistently showed the expression of just two ORs per sensilla, the many additional palpal ORs forwarded by Hodges et al. would suggest that most or all of the ~120 palpal ORNs would be co-expressing several tuning ORs, thus requiring a drastic revision of the neurobiological understanding of insect ORNs. Finally, even if we were to accept this view and assume that palpal sensilla do indeed co-express combinations of the novel tuning ORs described by Hodges et al., then it is reasonable to expect that electrophysiological responses to odorant stimuli would vary between the individual sensilla maxillary palp. However, this is in direct opposition to the uniformity of response between sensilla observed in multiple studies [2, 7, 10], and ignores the finding that in *An coluzzii*, the pharmacologic response profiles of AgOR8 and AgOR28 sufficiently explain the OR-mediated response profiles of the palpal sensilla [2, 7, 8, 11, 18, 19].

Instead, the simplest explanation for the surprising results reported by Hodges et al. is that during the gathering of their data sets they have—by any number of possible routes— inadvertently mixed antennal RNA in with that of the maxillary palp. To demonstrate the plausibility of this more expedient explanation, we compared the data from Hodges et al. to that of four other RNAseq data sets derived from the antennae and maxillary palps from three mosquito species and recently published by three independent research groups [7, 8, 11, 19].

The results of these comparisons as they pertain to ORs are presented in Figure 1 and Table 1. The first three panels of Figure 1 (A-C) show the relative abundances of OR transcript (reported in RPKM) in the maxillary palp for each of the annotated ORs in *An. coluzzii* and in *Ae. aegypti;* we have also included the three CO_2_ sensing GRs as additional points of reference. One can immediately appreciate the differences between the first two data sets (Figure 1A and 1B) and the data from Hodges et al. (Figure 1C). Previous transcriptome profiles of the maxillary palp of *An. coluzzii* (Figure 1A) and *Ae. aegypti* (Figure1B) consistently reveal high transcript levels for two canonical tuning ORs, Orco, and the three CO_2_ sensing GRs, with each of the GRs showing a transcript level similar to that of Orco. This same pattern was also seen in the transcriptome profile of the maxillary palp of the non-blood feeding mosquito, *Toxorhynchites amboinensis* [11] thus suggesting a stereotypical pattern of OR and GR expression in the maxillary palp that is conserved across all mosquito taxa. In contrast, Hodges et al. reported that 53 tuning ORs were expressed above 1 RPKM in the maxillary palp of *An. coluzzii*, with 26 being above 4 RPKM and 9 of them above the transcriptome-wide median level of 8.5 RPKM. While the two canonical palpal tuning ORs are among these ORs, they are **not** the most abundant tuning ORs in the palp, a startling observation that is contrary to other published accounts of OR expression in mosquito maxillary palp. The second relevant difference here—and a point that Hodges et al. do not address—is that while the expected three CO_2_ sensing GRs are indeed present in their maxillary palp data sets, the RPKM values for these GRs are more than an order of magnitude lower than values for the same GRs as seen in the maxillary palp of either *An. coluzzii* (Figure 1A) or *Ae. aegypti* (Figure 1B). While the ratios of transcript level between the canonical palpal GRs and tuning ORs appear similar across all these studies, the ratios of transcript level between these ORs and that of the OR co-receptor, Orco is highly divergent in the data from Hodges et al. So it appears that the palpal transcriptome profile reported by Hodges et al. contains a chemoreceptor profile that has some of the hallmarks of the other previous studies but that also appears to have been effectively diluted by a fraction of chemoreceptor-enriched, non-palpal RNA. We suggest that this fraction of non-palpal RNA is most likely derived from the very OR-rich antenna and has contaminated their maxillary palp samples.

**Table 1.** Rank order correlation (Spearman’s rho) of RPKM values for genes between antenna and maxillary palp.

| Study | Mosquito species | <u>Chemoreceptor Class</u> |  |  |  | <u>Transcriptome</u> |  |
| --- | --- | --- | --- | --- | --- | --- | --- |
|  |  | ORs | GRs | IRs | OBPs | Top 1000 genes in antenna** | T1000 antennal genes in Q3 of palp |
| Hodges et al. 2014<sup>†</sup> | *An. coluzzii* | **0.82**<br>*0.85\** | **0.79** | **0.95** | **0.95** | **0.76** | **2.7%** |
|  | *An. quadriannulatus* | **0.90**<br>*0.95* | **0.77** | **0.90** | **0.95** | **0.81** | **0.4%** |
| Pitts et al. 2011 | *An. coluzzii* | 0.13<br>*0.20* | 0.58 | 0.57 | 0.66 | 0.44 | 7.2% |
| Zhou et al. 2014 | *Ae. aegypti*<sup>‡</sup> | 0.05<br>*0.10* | 0.45 | 0.41 | 0.65 | 0.48 | 7.8% |
|  | *To. amboinensis* | 0.49<br>*0.58* | 0.52 | 0.48 | 0.55 | 0.49 | 5.8% |

<sup>†</sup> RPKM values analyzed and presented here were those reported by Hodges *et al.* in their supplementary materials.
<sup>‡</sup> Analysis based upon reanalyzed RNAseq data generated by two, unaffiliated and independent research groups that were published as part of two separate studies (Bohbot *et al.* 2013, McBride *et al.* 2014)
\*\*Pearson's R calculated on log-transformed RPKM values

**Figure 1.**
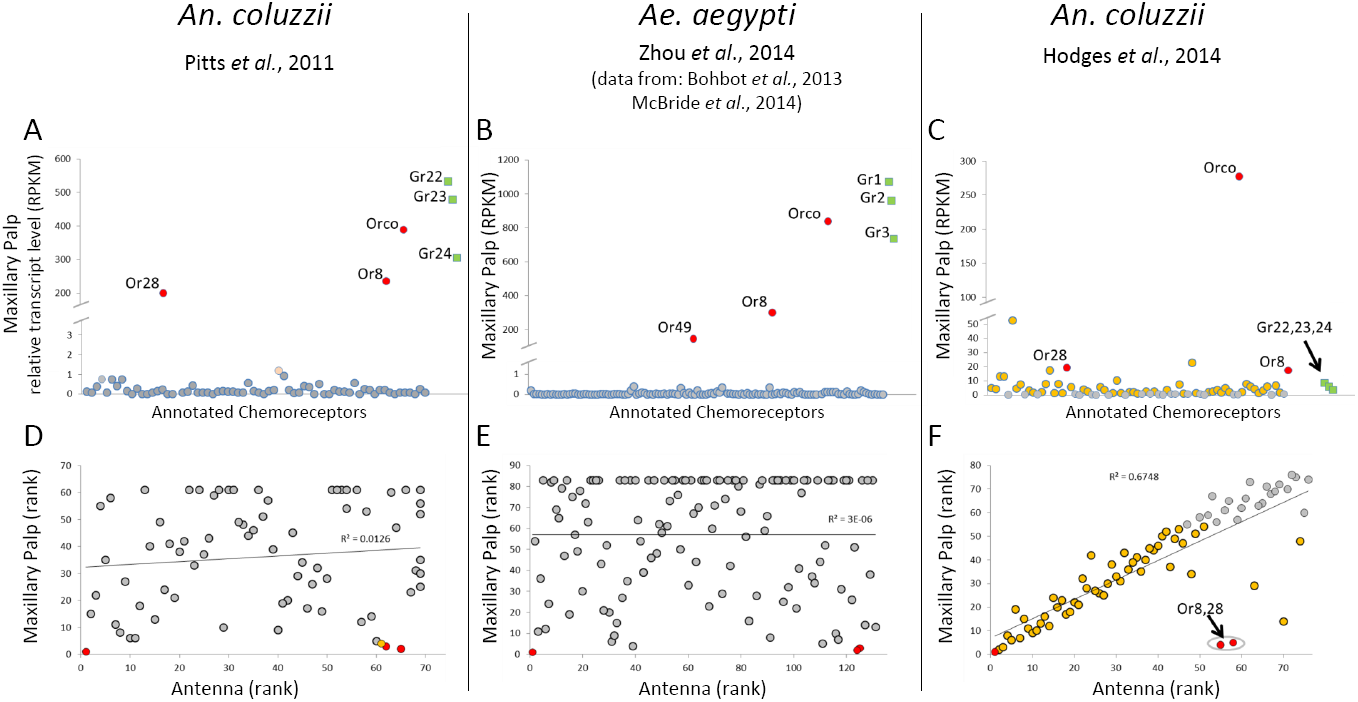
Expression levels of chemoreceptors in the chemosensory appendages of two mosquito species. Data are derived from RNAseq studies initiated by four, independent research groups as indicated below the species name. **A-C**: Internally normalized transcript levels (RPKM) of all the ORs (o) and the three CO2-sensing GRs (□) in the maxillary palp of the indicated mosquito species. **D-F**: Linear regression of the rank order (rank based upon RPKM) of the ORs in the antenna (y-axis) and the maxillary palp (x-axis). In all panels, ORs appearing above a base line of 1 RPKM in maxillary palp are highlighted in **orange**. The three canonical palpal ORs are indicated in **red**.

The possibility of antennal RNA contamination in the maxillary palp samples of Hodges et al. is further suggested by a pairwise comparison of the ranking (based upon RPKM level) of all the individual ORs between the maxillary palps and the antennae (Figure 1D-F). In the first two panels the rank orders of the antennal and palpal ORs from previously published reports in *An. coluzzii* (Figure1D) and *Ae. aegypti* (Figure 1E) show little to no correlation between the two tissues, consistent with the differences in the morphology and electrophysiology of the two appendages; this was also the case in *To. amboinensis* (Table 1). In contrast, the rank order of ORs in the palpal and antennal samples from Hodges et al. is very highly correlated (Figure 1F) with the most highly expressed ORs in the maxillary palp (orange points) also clustering among the most highly expressed ORs in the antenna. Importantly, the correlation shown in Figure 1F increases substantially once the two canonical palpal tuning ORs are removed from the analysis; a similar and even stronger trend is seen in the Hodges et al. palpal samples from *An. quadriannulatus* (Table 1). Furthermore, when we look to the expression profiles of other classes of chemosensory genes, the data from Hodges et al. continue to show correlations between antenna and maxillary palp that are consistently greater than those of other studies (Table 1). Even when we consider how well the ranks of the top 1000 genes in the antennal samples from Hodges et al. correlate with their ranking in the palpal samples, we see correlations that were absent in other mosquito studies (Table 1). Surprisingly, Hodges et al. present correlations among the chemoreceptor genes (Hodges et al., Figures 5 and 6) that are nearly identical to those shown in Figure 1F and Table 1, yet never address why it should be plausible that two developmentally-, morphologically-, neurologically-, and physiologically-distinct olfactory tissues should nevertheless look so identical in their patterns of chemoreceptor expression.

Hodges et al. suggest that their divergent findings regarding the expanded OR repertoire of the maxillary palps is related to the fact that their tissue samples were collected one hour following the onset of the mosquitoes’ dark phase (ZT13), whereas our previous analysis [8] used maxillary palps collected at a point prior to the onset of the dark phase (ZT10). To directly address the plausibility of this point, we preformed our own RNAseq analysis of *An. coluzzii* maxillary palp at precisely the same time point as Hodges et al, collecting tissues at the end of the first hour of the dark phase. The transcriptome profile from our new, ZT13 sample corresponded with our previous ZT10 palpal sample (Pearson’s R=0.95) and there was no indication of any of the many chemoreceptor enrichments reported by Hodges et al.

In summary we posit that the study by Hodges et al. is fundamentally compromised by the incorporation of contaminated RNA samples and the conclusions drawn are critically flawed. Accordingly, we strongly caution members of the scientific community who may wish to use the RNAseq data sets of Hodges et al. that doing so may lead to erroneous conclusions.

